# Charophytic Green Algae encode ancestral Pol IV/Pol V subunits and a CLSY/DRD1 homolog

**DOI:** 10.1101/2023.06.13.544724

**Authors:** Tania Chakraborty, Joshua T. Trujillo, Timmy Kendall, Rebecca A. Mosher

**Author notes:** These authors contributed equally (Tania Chakraborty, Joshua T. Trujillo).

## Abstract

In flowering plants, euchromatic transposons are transcriptionally silenced by RNA-directed DNA methylation (RdDM), a small RNA-guided *de novo* methylation pathway. RdDM requires the activity of the RNA Polymerase (Pol) IV and V, which produce small RNA precursors and non-coding targets of small RNAs, respectively. These polymerases are distinguished from Pol II by multiple plant-specific paralogous subunits. Most RdDM components are present in all land plants, and some have been found in the Charophytic green algae (CGA), a paraphyletic group that is sister to land plants. However, the evolutionary origin of key RdDM components, including the two largest subunits of Pol IV and Pol V, remains unclear. Here we show that multiple lineages of CGA encode a single-copy precursor of the largest subunits of Pol IV and Pol V, resolving the two presumed duplications in this gene family. We further demonstrate the presence of a Pol V-like C-terminal domain, suggesting that the earliest form of RdDM utilized a single Pol V-like polymerase. Finally, we reveal that CGAs encode a single CLSY/DRD1-type chromatin remodeling protein, further supporting the presence of a single specialized polymerase in CGA RdDM.

## Introduction

Eukaryotic genomes encode three highly conserved nuclear DNA-dependent RNA polymerases (Pol), each responsible for transcription of different subsets of RNA within the cell (Cramer *et al*., 2008; Werner & Grohmann, 2011). Transcription of ribosomal (r)RNA, protein-coding messenger (m)RNA, and transfer (t)RNA is regulated by Pol I, II, and III, respectively. These RNA Pols are holoenzyme complexes consisting of two large catalytic subunits and additional smaller non-catalytic subunits that regulate transcriptional activity and RNA processing. The largest two subunits of the complex possess highly conserved amino acid residues that constitute the active site, template DNA binding, and DNA-RNA hybrid binding regions. Each RNA Pol complex is composed of a unique pair of these catalytic subunits but many of the smaller subunits are shared between complexes. The evolution of distinct catalytic and accessory subunits has driven divergence in function between eukaryotic RNA Pol complexes.

Land plants are unique among eukaryotes in encoding two additional RNA polymerases, Pol IV and V, which are specialized for production of small interfering (si)RNA and longer non-coding transcripts in the RNA-directed DNA methylation (RdDM) pathway, respectively (Ream *et al*., 2014). RdDM is functionally similar to small RNA-mediated transcriptional gene silencing in fission yeast, utilizing an RNA-dependent RNA polymerase, a Dicer endonuclease, and an Argonaute protein to establish DNA methylation *de novo* (Matzke & Mosher, 2014). However, transcriptional silencing in fission yeast uses Pol II for both production of small RNAs and synthesis of non-coding transcripts that recruit the silencing machinery, while these activities are performed by Pol IV and Pol V in plants.

Pol II, IV, and V have 12 subunits that are designated <u>N</u>uclear <u>R</u>NA <u>P</u>olymerase (NRPx#) with x = B, D and E, respectively, and # = 1-12 (largest to smallest subunit) (Ream *et al*., 2014). Although most subunits are shared between the three complexes, some are shared between two of the three while others are unique to a single complex. Pol IV and V are primarily distinguished from Pol II by unique largest subunits, NRPD1 and NRPE1, respectively. Furthermore, Pol IV and V share a second-largest subunit (NRPD/E2), which is distinct from the Pol II second-largest subunit (NRPB2). The catalytic domains of NRPD1, NRPE1, and NRPD/E2 have diverged from NRPB1, but key amino acids and domains associated with transcription are conserved (Ream *et al*., 2014).

Although clearly homologous in the catalytic region, the largest subunits are significantly diverged at the carboxyl terminal domain that serves as a binding platform for interacting proteins (Ream *et al*., 2014). NRPB1 contains well-characterized seven amino acid (heptad) repeats while NRPD1 and NRPE1 instead possess a domain of unknown function (DUF3223 or DeCL domain) (Ream *et al*., 2014). NRPE1 also contains an intrinsically disordered, repetitive, and evolutionarily labile region between the catalytic and DeCL domains (Trujillo *et al*., 2016). This region is enriched in AGO hooks, peptide motifs that mediate interaction with Argonaute proteins (El-Shami *et al*., 2007).

Pol IV and Pol V also differ in how they are recruited to DNA. Pol IV requires CLSY proteins, which are part of the SNF2 ATPase family of chromatin remodelers. Arabidopsis possesses four CLSY proteins, which recruit Pol IV in a tissue and locus-specific fashion (Zhou *et al*., 2018). Pol V instead requires DRD1, the only other member of the CLSY subclade of SNF2 ATPases, for association with DNA (Zhong *et al*., 2012). While evolutionary histories of other RdDM components have been elucidated, we lack information on the evolutionary origin of CLSY/DRD1 proteins.

Previous studies indicate that Pol IV and V arose prior to the emergence of land plants through the retention of paralogous subunits following the duplication of RNA Pol II subunits (Luo & Hall, 2007). Land plants are sister to green algae and together form the Viridiplantae (green plants) lineage, all of which share a common ancestor (Becker & Marin, 2009). Green algae consist of several different lineages that are divided among the monophyletic clade of Chlorophytic green (Chlorophyta) and a polyphyletic sister group called Charophytic green algae (CGA). Together CGA and land plants form a monophyletic clade known as Streptophyta (de Vries & Archibald, 2018) (**Supplemental Figure S1**). Although Pol IV and V subunits are absent in Chlorophytic green algae, a putative NRPD1 homolog was identified in a single CGA order, suggesting a Pol IV-like complex evolved immediately prior to the emergence of land plants (Luo & Hall, 2007). However, the origin of this homolog, other Pol IV and V subunits, and Pol-associated RdDM components remains unclear.

Here, we report that a single NRPD1/NRPE1-like sequence is present in multiple CGA lineages. This sequence has many of the structural hallmarks of NRPD1 and NRPE1, but has a CTD similar to Pol V. In addition, we demonstrate that a single CLSY/DRD1 protein is present in the CGAs, raising the possibility that a primitive RdDM utilizing a single specialized polymerase evolved in plants prior to terrestrialization.

## Results

### CGA taxa encode a single NRPD1/NRPE1 homolog, and it is expressed in Penium margaritaceum

To examine whether Charophytic Green Algae (CGA) possess homologs of *NRPD1* and *NRPE1* we queried the protein and transcript databases of selected CGA taxa by using the *Arabidopsis thaliana* NRPB1, NRPD1, and NRPE1 sequences. We assessed genomes of both early- and late-diverging CGA taxa, including Klebsormidiophyceae, Charophyceae, and Zygnematophyceae. Protein BLAST identified NRPB1 homologs in multiple lineages of CGA. These genes were generally well annotated and supported by transcriptome data. In contrast, similar searches for NRPD1 and NRPE1 obtained partial gene predictions that matched mostly to the catalytic region of the protein, where conservation between NRPB1, NRPD1, and NRPE1 is highest (Kanno *et al*., 2005; Pontier *et al*., 2005; Herr, 2005; Luo & Hall, 2007). We identified partial transcripts that share homology with Arabidopsis NRPD1 and NRPE1 in *Chlorokybus atmophyticus, Klebsormidium nitens, Entransia fimbriata, Chara braunii, Coleochaete irregularis, Spirogya* sp., *Mougeotia scalaris, Mesotaenium kramstae*, and *Penium margaritaceum* (**Table S1**). To further investigate these predicted transcripts, we amplified partial cDNA sequences from *Penium margaritaceum* and sequenced the resulting fragments. The sequenced cDNA fragments have numerous differences with respect to the predicted transcript, but show high similarity to Arabidopsis NRPD1 and NRPE1 (**Supplemental Dataset 1**).

To examine the evolutionary relationship between known Pol IV and Pol V subunits and the predicted first subunit homologs in CGA taxa, we constructed a multi-sequence alignment and inferred phylogeny using the Maximum Likelihood method (**Figure 1**). Two major clades consist of NRPB1 and NRPD1/NRPE1 homologs from across green plants, with the newly identified CGA homologs diverging prior to the division of land plant NRPD1 and NRPE1. This position indicates that the predicted CGA sequences represent the single-copy ancestor of NRPD1 and NRPE1. Our result agrees with the 2007 Luo and Hall study, and their predicted sequences fall in the same intermediate position as our predicted CGA first subunits. To confirm that the predicted CGA first subunit homologs are not homologs of Pol I or Pol III largest subunits, we also inferred phylogeny from a multi-sequence alignment containing NRPA1 and NRPC1 homologs. We observed that the NRPA1 and NRPC1 sequences form distinct clades, and the CGA NRPD/E1 sequences remain in the intermediate position between the NRPB1 clade and the land plant NRPD1 and NRPE1 clades (**Supplemental Figure S2**).

**Figure 1.**
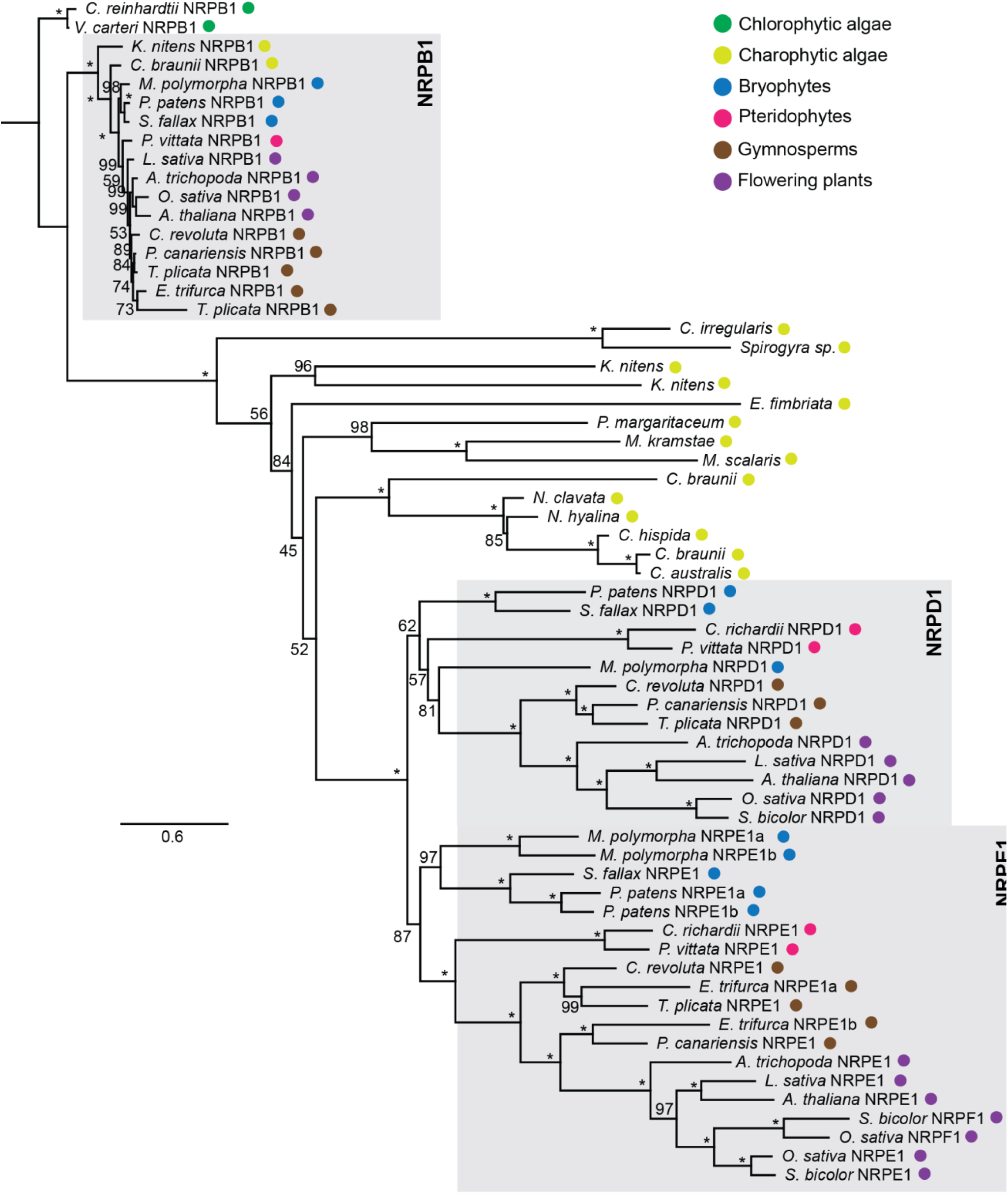
Phylogenetic analysis of NRPB1, NRPE1, and NRPD1 homologs shows a Pol IV/Pol V-like first subunit in multiple Charophytic green algae. Amino acid sequences from conserved catalytic regions (C-G domains) were aligned with MAFFT v7.450 and stripped in positions where 50% of the taxa contained a gap. The tree was inferred by maximum likelihood and rooted on Chlorophyte NRPB1 sequences. Bootstrap support is listed on each brand (*, 100% support).

### CGA first subunit homologs resemble NRPD1 and NRPE1 at key catalytic motifs

NRPD1 and NRPE1 sequences have a higher rate of substitution than NRPB1 sequences, resulting in longer branch lengths (Luo & Hall, 2007). Despite this sequence divergence, the Metal A site in domain D remains conserved across NRPB1, NRPD1, and NRPE1 (Rymen *et al*., 2020). The Metal A site, which binds a magnesium ion near the polymerase active site, is essential for Pol IV and Pol V function, as substituting the aspartic acid residues in Arabidopsis NRPD1 or NRPE1 causes depletion of siRNA production and loss of DNA methylation (Haag *et al*., 2009). To examine whether the CGA first subunit homologs also possess the Metal A site, we searched the multi-sequence alignment and found the “DFDGD” sequence which represents the core of the Metal A site in all the CGA first subunits (**Figure 2**). The strict conservation of these five residues within the CGA first subunit homologs suggests that they could assemble into a catalytically active polymerase holoenzyme. Haag et al. noted that the broader conservation pattern around the DFDGD residues is not maintained in land plant NRPD1 and NRPE1 sequences, a pattern that extends into the CGA first subunits we evaluated (**Figure 2**).

**Figure 2.**
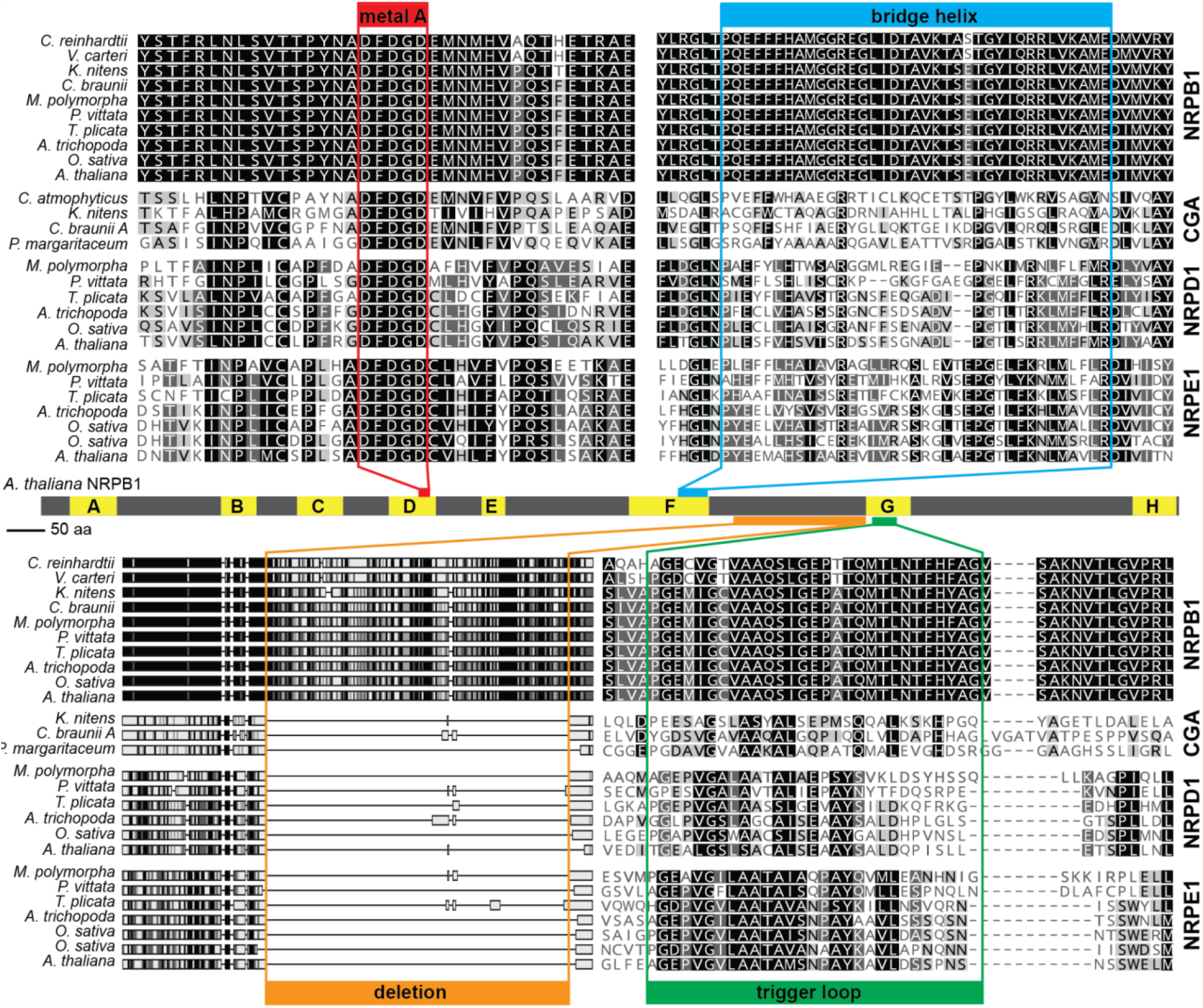
CGA homologs show hallmarks of NRPD1 and NRPE1 in the catalytic regio. Alignment of NRPB1, NRPD1, and NRPE1 sequences from land plants and CGAs demonstrates conservation of the Metal A site, loss of conservation in the bridge helix and trigger loop, and a large deletion between domains F and G. All amino acid sequences were aligned with MUSCLE; sequences from a single gene were extracted from the alignment and shaded by similarity in Geneious Prime.

Unlike the Metal A site, there is significant divergence between NRPB1 and NRPD1/NRPE1 sequences in other functionally important motifs. The bridge helix domain is important for NRPB1 catalytic activity, and the 35-residue region is highly conserved in NRPB1 sequences across the different kingdoms of life (Weinzierl, 2010; Hein & Landick, 2010). However, conservation in this region is lost in NRPD1 and NRPE1 (Herr, 2005). We examined the bridge helix region in the CGA first subunit homologs and observed substantial divergence from the NRPB1 sequence (**Figure 2**).

NRPD1 and NRPE1 homologs also possess substitutions and deletions relative to NRPB1 within the trigger loop region (Ferrafiat *et al*., 2019; Rymen *et al*., 2020). Loss of the trigger loop causes errors in Pol II transcription, and the loss of conservation of this region might explain Pol IV’s error-prone transcription (Kaplan *et al*., 2008; Erhard *et al*., 2009; Haag *et al*., 2012; Marasco *et al*., 2017; Rymen *et al*., 2020). We examined the trigger loop in CGA largest subunit homologs and observed that it is not conserved in the CGA sequences, making them look more like NRPD1 and NRPE1 than NRPB1 (**Figure 2**). Immediately upstream of the trigger loop, NRPD1 and NRPE1 also have a ∼190 amino acid deletion of the foot domain (Luo & Hall, 2007; Matzke *et al*., 2015). We confirmed that this deletion is conserved across land plant lineages and also exists in the CGA homologs (**Figure 2**). Together, analysis of domains and motifs in the CGA largest subunit homologs shows that these sequences contain the sequence features that are characteristic of Pol IV and Pol V largest subunits, further supporting the phylogenetic placement of these CGA sequences as single-copy ancestors of NRPD1 and NRPE1.

### The CGA NRPD1/NRPE1 homologs have C-terminal domains with AGO hooks

Although NRPD1 and NRPE1 are clearly homologous to NRPB1 throughout their catalytic regions, their carboxy-terminal domains (CTD) are not homologous and likely arose through a gene fusion event (Ream *et al*., 2014). The NRPB1 CTD consists of multiple repeats of seven amino acids (YTPTSPS), a motif that is conserved across all eukaryotic NRPB1s. The CTDs of NRPD1 and NRPE1 lack this motif and instead possess a domain related to the Defective in Chloroplasts and Leaves (DeCL) protein. NRPE1 also has an AGO-binding platform, an intrinsically disordered and repetitive region enriched in “AGO hooks” (GW, WG, or GWG peptides) (Till *et al*., 2007; El-Shami *et al*., 2007; Trujillo *et al*., 2016). Luo and Hall identified a single additional polymerase subunit in members of the Charales and they designated these sequences as (N)RPD1, in part due to their conclusion that NRPE1 sequences were absent from non-flowering plants (Luo & Hall, 2007). Subsequently, NRPE1 sequences were identified in all lineages of land plants, raising the possibility that the single additional subunit in CGA lineages might be more similar to NRPE1 in the C-terminal region, or even retain the heptad repeats found in NRPB1. To examine whether the CGA largest subunit homologs more closely resemble NRPB1, NRPD1, or NRPE1 in the CTDs, we searched for AGO hook motifs and the DeCL domain in regions downstream of the final catalytic domain. We identified numerous AGO hooks in the *K. nitens, C. braunii*, and *P. margaritaceum* homologs, as well as predicted DeCL domains in *K. nitens* and *C. braunii* (**Figure 3A**). We confirmed the presence of AGO hooks in the *P. margaritaceum* transcript through RT-PCR and recovered multiple alleles at the C-terminus (**Supplemental Dataset 1**). However, our *P. margaritaceum* cDNA fragments do not contain stop codons and therefore may not contain the extreme C-terminus. The presence of AGO hooks and a DeCL domain in these homologs indicates that the additional NRPB1 homologs in CGAs more closely resemble NRPD1 than NRPE1, and probably interact with AGO proteins.

**Figure 3.**
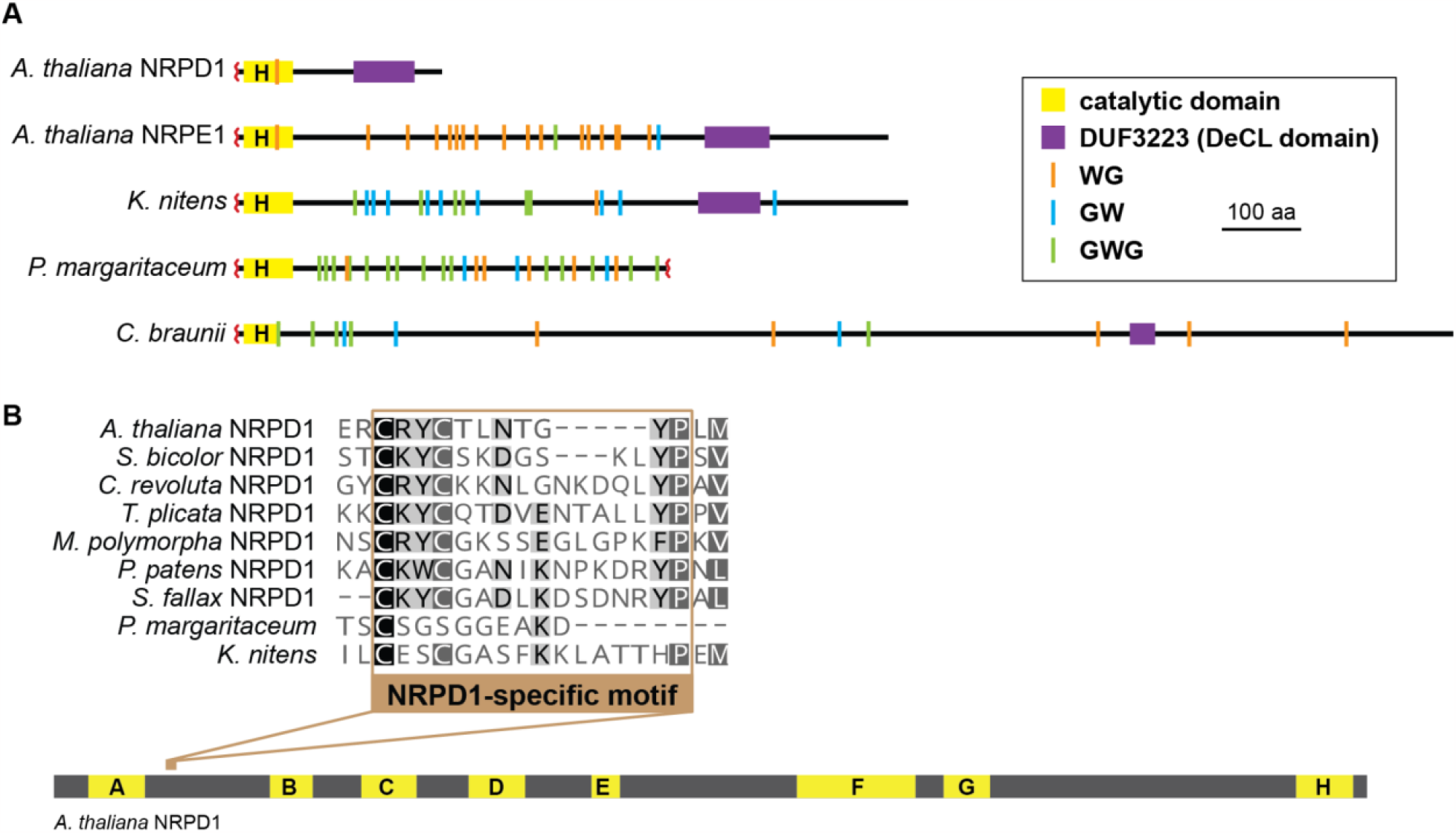
CGA homologs resemble NRPE1 in the C-terminal domain. (**A**) Diagram of the carboxy-terminal domain (CTD) of *Arabidopsis* NRPD1, *Arabidopsis* NRPE1, and homologous sequences from *K. nitens, P. margaritaceum*, and *C. braunii* demonstrating the presence of numerous AGO hooks (WG, GW, or GWG peptides) between the catalytic region and the DeCL domain. (**B**) Alignment of the NRPD1-specific motif (Ferrafiat et al., 2019) from land plant NRPD1 and homologous CGA sequences demonstrates conservation of this motif across land plants is absent in the CGA sequences.

Mutations in NRPD1-specific motif (C[KR]YC) reduce siRNA accumulation, with mutations in the cysteine 118 residue being particularly detrimental (Ferrafiat *et al*., 2019). The NRPD1-specific motif is conserved in angiosperms and at least two gymnosperms (Ferrafiat *et al*., 2019). To examine whether non-seed plant NRPD1 and the CGA NRPD/E1 homologs possess this motif, we examined the alignment of conserved A-H domains across selected land plant and CGA species. We observed that the C[KR]YC box is conserved across land plants, including bryophytes, but this sequence is not found in the CGA homologs (**Figure 3B**). This observation further indicates that the additional largest subunit homologs in CGA more closely resemble NRPE1 than NRPD1.

### Later-diverging CGA taxa encode an NRPD/E2 homolog

Pol IV and Pol V share a second subunit, NRPD/E2, which is conserved across all land plants but was not previously identified in CGA species (Luo & Hall, 2007; Huang *et al*., 2015; Wang & Ma, 2015). To examine whether the CGAs possess non-canonical second subunit homologs, we utilized BLAST to identify second subunit homologs in CGA genomes. We identified NRPB2 homologs across all the land plant lineages as well as CGAs and Chlorophytic algae, and we identified NRPD/E2 homologs in the tested land plant genomes. We also observed additional second subunit sequences in *E. fimbricata, C. braunii, M. kramstae*, and *P. margaritaceum*, CGAs from the Zygnematales and Charales orders, suggesting that these later-diverging CGAs might contain a non-canonical second subunit (**Supplementary Table 2**). To identify if these sequences are NRPD/E2 homologs, we first inferred phylogeny from Maximum Likelihood analyses. The topology of these phylogenetic trees indicates that the partial sequences we identified are NRPD/E2-like (**Figure 4**). To examine whether the CGA homologous sequences are truly intermediate between the NRPB1 clade and the land plant NRPD/E2, we gathered NRPA2 and NRPC2 sequences across the land plant lineage as well as from CGA taxa and constructed a multi-sequence alignment with all the second subunit homologs. We observed that the CGA NRPD/E2 sequences remain outside the well-supported NRPA2 and NRPC2 clades (**Supplemental Figure S3**).

**Figure 4.**
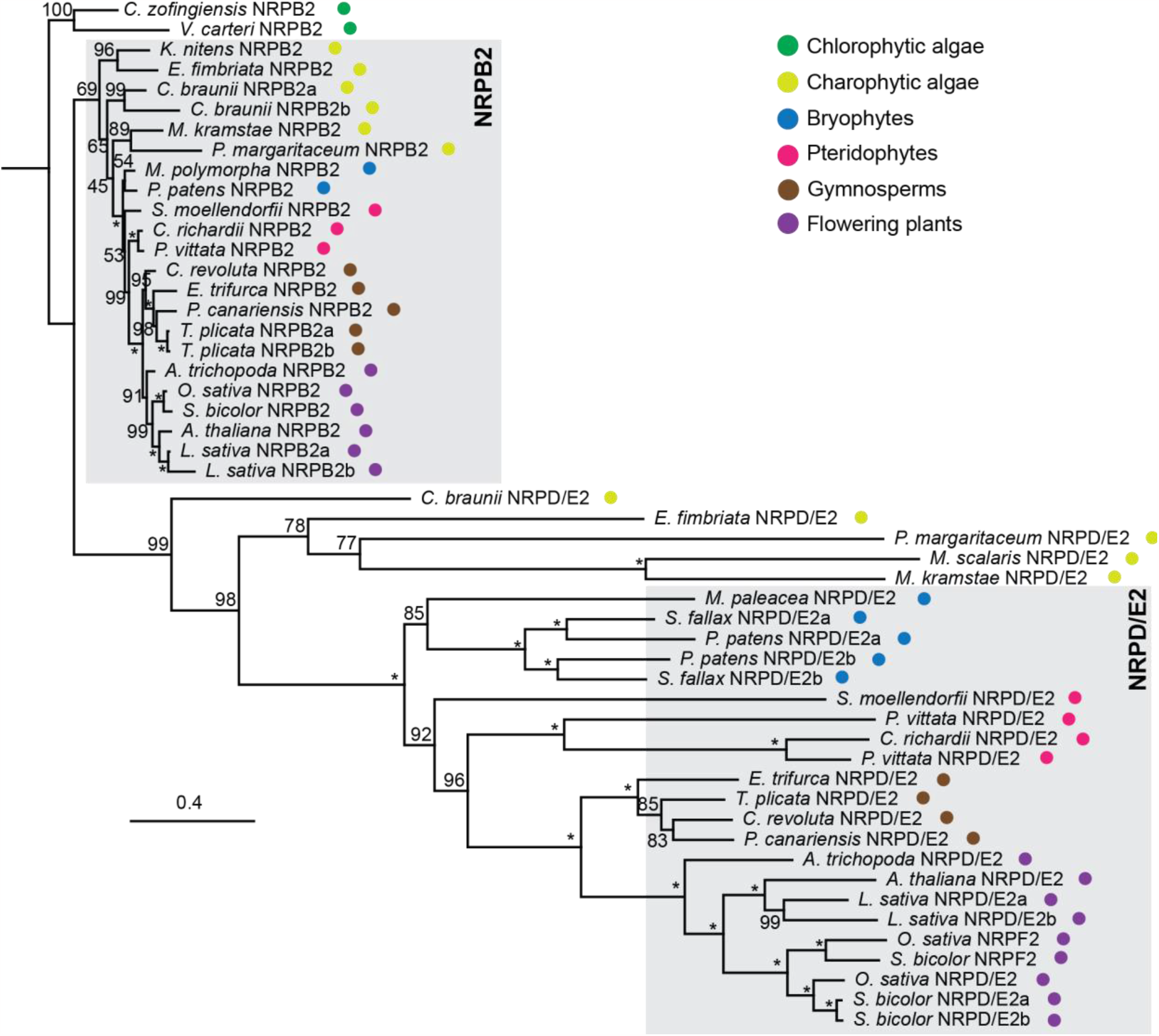
Phylogenetic analysis of NRPB2 and NRPD/E2 homologs shows a Pol IV/Pol V-like second subunit in multiple Charophytic green algae. Amino acid sequences were aligned with MAFFT v7.450, and stripped in positions where 80% of the taxa contained a gap. The tree was inferred by maximum likelihood and rooted on Chlorophyte NRPB2 sequences. Bootstrap support is listed on each brand (*, 100% support).

To further examine the identity of these sequences, we assessed characteristic functional domains in the second subunit. Similar to first subunit homologs, second subunits of RNA polymerases contain a metal binding site that is responsible for complexing magnesium in the active site. A multi-sequence alignment of second subunit sequences demonstrates high conservation of the metal B site across all sequences, indicating that the additional CGA second subunit homologs may be catalytically active polymerase subunits (**Figure 5**). The hybrid-binding site of NRPB2 binds the nascent DNA-RNA hybrid and interacts with the bridge helix (Cramer *et al*., 2001). Like the bridge helix, the hybrid-binding site in NRPD/E2 is less conserved than in NRPB2 across the land plant lineage (Herr, 2005). We observes that CGA second subunit homologs also have low conservation in this region (**Figure 5**).

**Figure 5.**
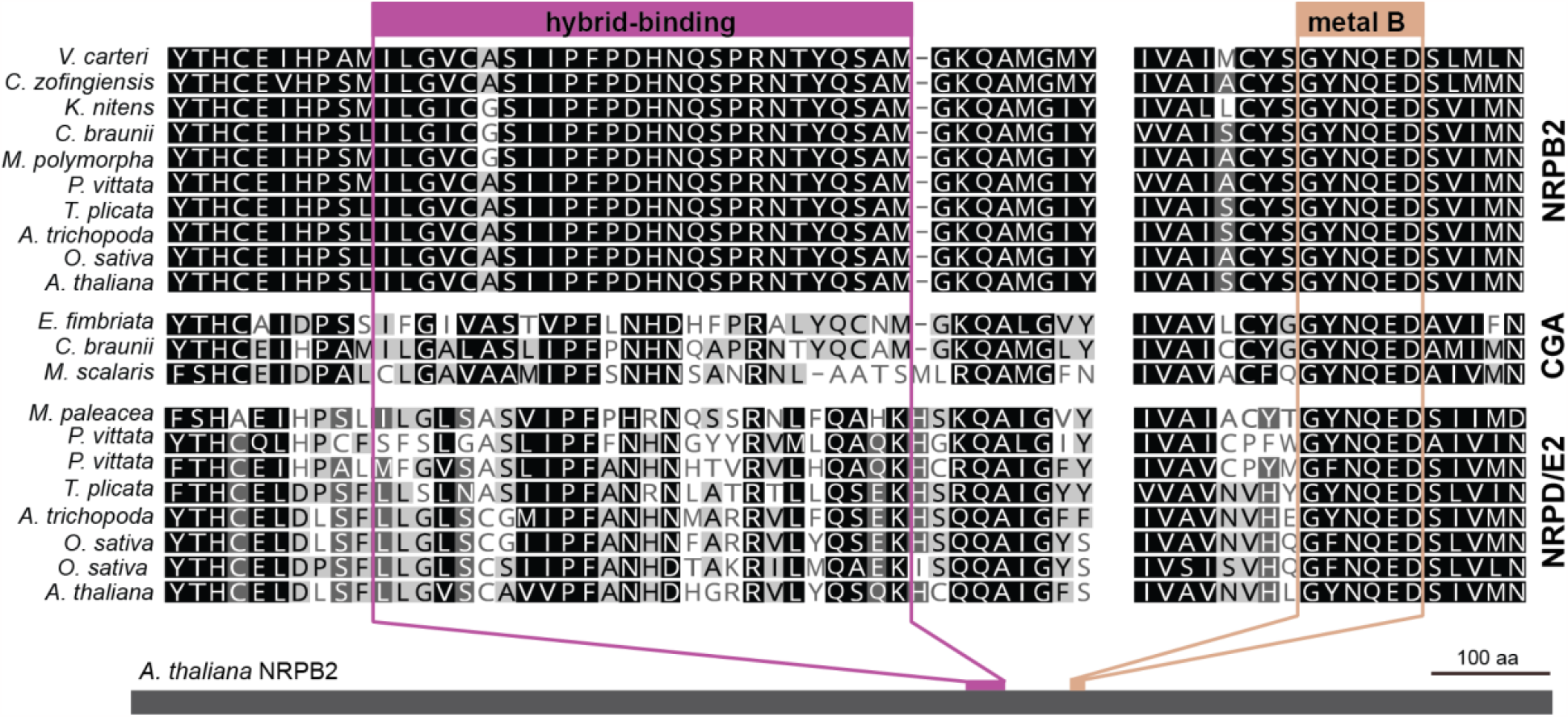
CGA homologs resemble NRPD/E2 at key motifs. Alignment of conserved domains of Pol II, Pol IV, Pol V second subunit sequences from land plants and CGAs demonstrates conservation of the Metal B site and lack of conservation in the hybrid-binding domain. All amino acid sequences were aligned with MUSCLE; sequences from a single gene were extracted from the alignment and shaded by similarity in Geneious Prime. The partial sequence from *Penium margaritaceum* did not contain these regions.

Together, these observations support the presence of an NRPD/E2 subunit in at least some CGA lineages. Taken together, our data indicate the first plant specific homolog of Pol II arose in the CGAs, prior to its duplication into separate Pol IV and Pol V enzymes in land plants.

### A single CLSY/DRD1 sequence is also present in CGAs and is expressed in Penium

Pol IV and V require SNF2 ATPase homologs of the CLSY/DRD1 family for their activity. Arabidopsis contains four CLSY proteins, and the quadruple mutant phenocopies Pol IV mutants with respect to siRNA production (Smith *et al*., 2007; Zhou *et al*., 2022). There are two additional homologs in this small gene family, DRD1 and CHR34 (Kanno *et al*., 2004; Knizewski *et al*., 2008). DRD1 is required for Pol V association to DNA (Zhong *et al*., 2012), and CHR34 has no known role in RdDM. Since CGAs encode Pol IV/Pol V-like first and second subunits, we hypothesized that CGAs would also encode homologs of the CLSY/DRD1 proteins. We used BLAST to gather peptide sequences sharing homology with the Arabidopsis CLSY/DRD family proteins across land plants and in the CGAs. We observed CLSY and DRD1 homologs across land plants in many CGA taxa. To examine whether this single CLSY/DRD1 protein is expressed, we sequenced cDNA fragments from *Klebsormidium nitens* and *Penium margaritaceum* (**Supplemental Dataset 1**). The single CLSY/DRD1 was expressed in both species.

To determine the evolutionary relationship between the CGA CLSY/DRD1 homologs, we inferred a phylogeny that included land plant CLSY and DRD1 homologs as well as homologs of RAD54 and ATRX, the next most closely-related SNF2 family members (Knizewski *et al*., 2008). We observed three pairs of related proteins in Arabidopsis, representing DRD1/CHR34, CLSY1/CLSY2, and CLSY3/CLSY4. In most vascular plant, these protein pairs exist as single homologs (two CLSY-type, and one DRD1), while bryophytes encode a single CLSY homolog and a single DRD1 (**Figure 6** and **Supplementary Figure S4**). CGA genomes reduce this complexity further, encoding only one CLSY/DRD1-like protein, which are positioned sister to the land plant DRD1 and CLSY clades (**Figure 6**). This pattern suggests that CGAs, which encode a single Pol IV/V-like polymerase, also encode a single CLSY/DRD1 homolog. The CLSY and DRD1 proteins diverged in the ancestor of bryophytes coincident with the divergence of Pol IV and Pol V.

**Figure 6.**
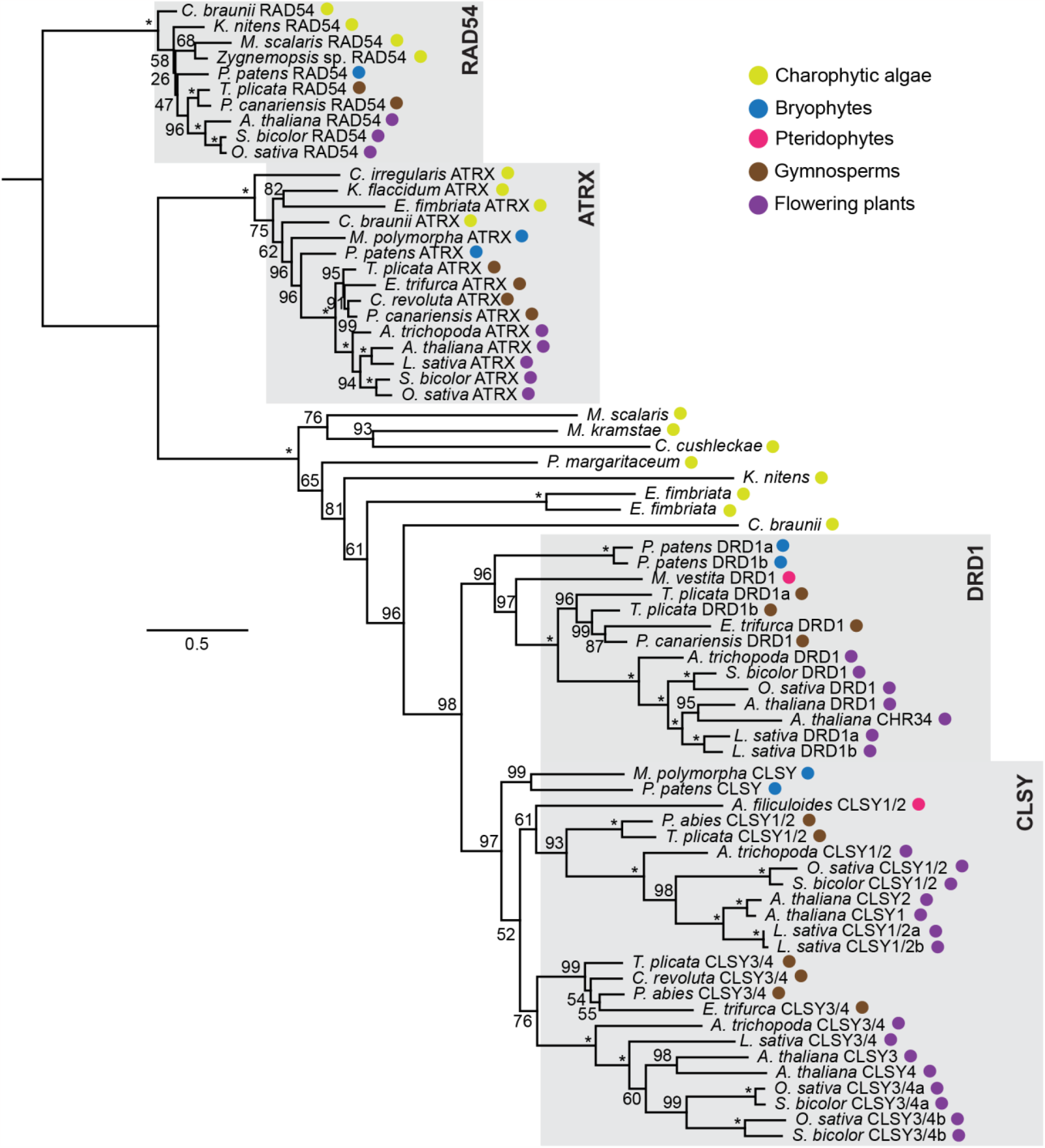
Phylogenetic analysis of CLSY and DRD1 homologs identifies a single homolog in multiple Charophytic green algae. Amino acid sequences were aligned with MAFFT v7.450, the conserved SNF2 and helicase domains and intervening sequences were extracted, realigned, and stripped in positions where 50% of the taxa contained a gap. The tree was inferred by maximum likelihood and rooted on RAD54 sequences. Bootstrap support is listed on each brand (*, 100% support).

## Discussion

Duplication and diversification of multiple Pol II subunits has led to the presence of Pol IV and Pol V in all land plants lineages (Huang *et al*., 2015; Wang & Ma, 2015). These specialized polymerases produce RNAs responsible for RNA-directed DNA Methylation (Wendte & Pikaard, 2017). Here we demonstrate that Charophytic green algae, the sister lineages to land plants, contain largest and second-largest subunits related to Pol IV and Pol V, as well as a homolog of CLSY/DRD1, which is required for Pol IV and Pol V activity. Together with observation of other RdDM components in CGAs, our results suggest that CGAs might perform an ancient form of RNA-directed DNA methylation using only a single additional polymerase.

An NRPD1-like transcript was previously identified in two genera within Charales, which at the time was considered the lineage most closely related to land plants (Luo & Hall, 2007). However, this studied failed to identify NRPD1-like sequences in other CGA orders and found no evidence for NRPD/E2-like sequences in CGAs. Our study identifies first subunit homologs in the four CGA orders most closely related to land plants (Klebsormidiales, Charales, Coleochaetales, and Zygnematales, **Figure 1**). We also identified a partial transcript in *Chlorokybus atmophyticus* (Chlorokybales order) with similarity to Pol IV and Pol V largest subunits, hinting that all CGA orders might contain a specialized polymerase (**Supplemental Table S1, Supplemental Figure S2**). We identified a second subunit homolog in Charales, Zygnematales, and Klebsormidiales, but not in earlier diverging orders (**Figure 4, Supplemental Figure S3**), suggesting asynchronous evolution of the largest two subunits. Similarly, we were unable to identify paralogs of the Pol IV and Pol V seventh subunit in any CGA genome, despite the presence of this subunit in all land plants (Huang *et al*., 2015). These observations are in line with presumed increasing elaboration of Pol IV and Pol V holoenzyme assemblies over evolutionary time (Luo & Hall, 2007; Huang *et al*., 2015).

Analysis of the conserved functional motifs in the first and second subunits reveals how closely the CGA subunits resemble land plant Pol IV and Pol V, suggesting that these subunits might function together in a single polymerase (**Figures 2 and 5**). Biochemical analysis is necessary to demonstrate such an interaction and to determine whether the resulting polymerase has the enzymatic characteristics of RNA Pol IV or Pol V (Marasco *et al*., 2017). However, the C-terminal domain reveals that CGA first subunit homologs are more similar to NRPE1, suggesting that a polymerase formed from this subunit might function like Pol V (**Figure 3**).

Our analyses also shows that a single CLSY/DRD1 homolog is present in CGA lineages (**Figure 6**), paralleling the presence of a single specialized largest polymerase subunit. Other components of RdDM, including RDR2, DCL3, AGO4, and DRM2, also have paralogous sequences in CGA genomes (You *et al*., 2017; de Mendoza *et al*., 2018; Wang *et al*., 2021; Bélanger *et al*., 2023), raising the possibility that CGAs perform some form of RdDM. The presence of a single specialized polymerase with a Pol V-like tail suggests two models for RdDM in CGAs. First, it is possible that the specialized CGA polymerase produces both siRNA precursors (like Pol IV) and the non-coding RNA scaffold responsible for recruiting siRNA-containing Argonaute complexes (like Pol V). These functions might then have been subfunctionalized into NRPD1 and NRPE1 in land plants. Alternatively, CGA RdDM might utilize Pol II for siRNAs production while its specialized polymerase functions like Pol V in land plants. Further research into this intriguing group of algae is needed to reveal the earliest forms of RdDM and expand our understanding of the diverse ways epigenomes are maintained.

## Materials and Methods

### Identification of homologs across green plant lineage

Homologs for all the proteins involved in the study were obtained in *Oryza sativa, Sorghum bicolor, Lactuca sativa*, and *Thuja plicata* by protein BLAST searches across land plants in Phytozome 12 and 13 using *Arabidopsis thaliana* peptide sequences as query. Sequences in other land plants were obtained either via protein BLAST in Phytozome 12 and 13 using *Arabidopsis thaliana* peptide sequences as query or from submitted data for Huang et. al 2015 (accessed from TreeBase, study 16473, last accessed on 3rd January, 2023). For *Cycas revoluta* first subunit sequences, we accessed the shotgun transcriptome data submitted by Huang et al. (EMBL GenBank ID GBJU00000000) and used the transcriptome for search using *Arabidopsis thaliana* peptides as queries. For Charophytic Green Algae, we utilized a broad range of search techniques. We obtained genomic, transcriptomic, and peptide databases for *Klebsormidium nitens, Chara braunii*, and *Mesotaenium kramstae* from Phycocosm (https://phycocosm.jgi.doe.gov/, (Hori *et al*., 2014; Nishiyama *et al*., 2018; Grigoriev *et al*., 2021)), and *Penium margaritaceum* sequence from the Penium genome database (Jiao *et al*., 2020). The *Chara braunii* genome is also hosted by OrcAE (http://bioinformatics.psb.ugent.be/orcae/overview/Chbra) and was utilized to search for additional sequences. These resources were searched by pBLAST using Arabidopsis peptide sequences as query. Peptide matches were evaluated by checking whether they contained the expected conserved domains. Transcriptome searches for CGAs were conducted by downloading 1kP transcriptomes from Cyverse (One Thousand Plant Transcriptomes Initiative, 2019; Carpenter *et al*., 2019) and using tblastn with Arabidopsis peptide sequences. Blast hits to genomic sequences were examined by taking 10-15 kb of nucleotide sequence around the blast hit/hits and finding predicted exons with fgenesh and fgenesh+ (Soloyev et al 2007) as an unbiased approach to exon finding, or by using the Arabidopsis peptide as reference respectively. ORFs underlying the predicted exons were defined and translated in their frame of reference for generating peptide sequences.

### Culturing of *P. margaritaceum* and *K. nitens*

*P. margaritaceum* (supplied by Professor Jocelyn Rose at Cornell University) and *K. nitens* (University of Texas at Austin Culture Collection, UTEX 623, *Klebsormidium flaccidum*) were cultured in agar slants containing Bristol medium (2.94 mM NaNO_3_, 0.17mM CaCl_2_.2H_2_0, 0.3 mM MgSO_4_.7H_2_O, 0.43 mM K_2_HPO_4_, 1.29 mM KH_2_PO_4_, 0.43 mM NaCl) supplemented with UTEX Soilwater: GR+ Medium under constant lighting at 25°C. The cultures were maintained by transferring once a month.

### RT-PCR and sequencing of Penium and Klebsormidium transcripts

Algal culture was scraped from the slant surface and total nucleic acid was extracted following a protocol adapted from White and Kaper (1989), with the modification of performing the phenol-chloroform-isoamyl alcohol extraction step twice. About 1.1-1.3 µg of total nucleic acid was subjected to DNAse treatment using Invitrogen DNA Free kit (catalog no. AM1906). 2.5 µg of DNAse-treated RNA was used in a 20 µL reaction to convert into cDNA using ThermoFisher SuperScript™ IV First-Strand Synthesis System (catalog number 18091050) using random hexamer primers. 1 µL of the reaction was then used to amplify predicted transcripts using gene-specific primers. The amplicon was allowed to run out on agarose gel, and the fragment was excised and purified using GeneJET Gel Extraction and DNA Cleanup Micro Kit (catalog number K0831). The fragments were T-cloned using the pGEM®-T Easy Vector System I (cat A1360, Promega) and plated on X-gal/IPTG plates. Three white colonies were selected to undergo restriction digestion with EcoR I to confirm transgene insertion, and upon confirmation, plasmids were sequenced at Plasmidsaurus to generate the cDNA sequence.

### Phylogenetic analysis

Multi-subunit alignments were constructed using MAFFT v7.450 on the Geneious Prime (version 2021.2, DotMatics) platform. Alignments were curated manually and gap trimming was performed using Mask Alignment option in Geneious on sites with gaps in at least 80% of the taxa. Phylogenetic inference was made using the iq-TREE web server using options LG for substitution model, +R free-rate heterogeneity, and Ultrafast Branch Support Analysis using 1000 bootstrap alignments and default Search Parameters (Nguyen *et al*., 2015; Hoang *et al*., 2018).

## Supplemental Material

**Supplemental Table S1: taxa for Figure 1**

**Supplemental Table S2: taxa for Figure 4**

**Supplemental Table S3: taxa for Figure 6**

**Supplemental Dataset 1: cDNA sequences from *Penium margaritaceum***

**Supplemental Figure S1.**
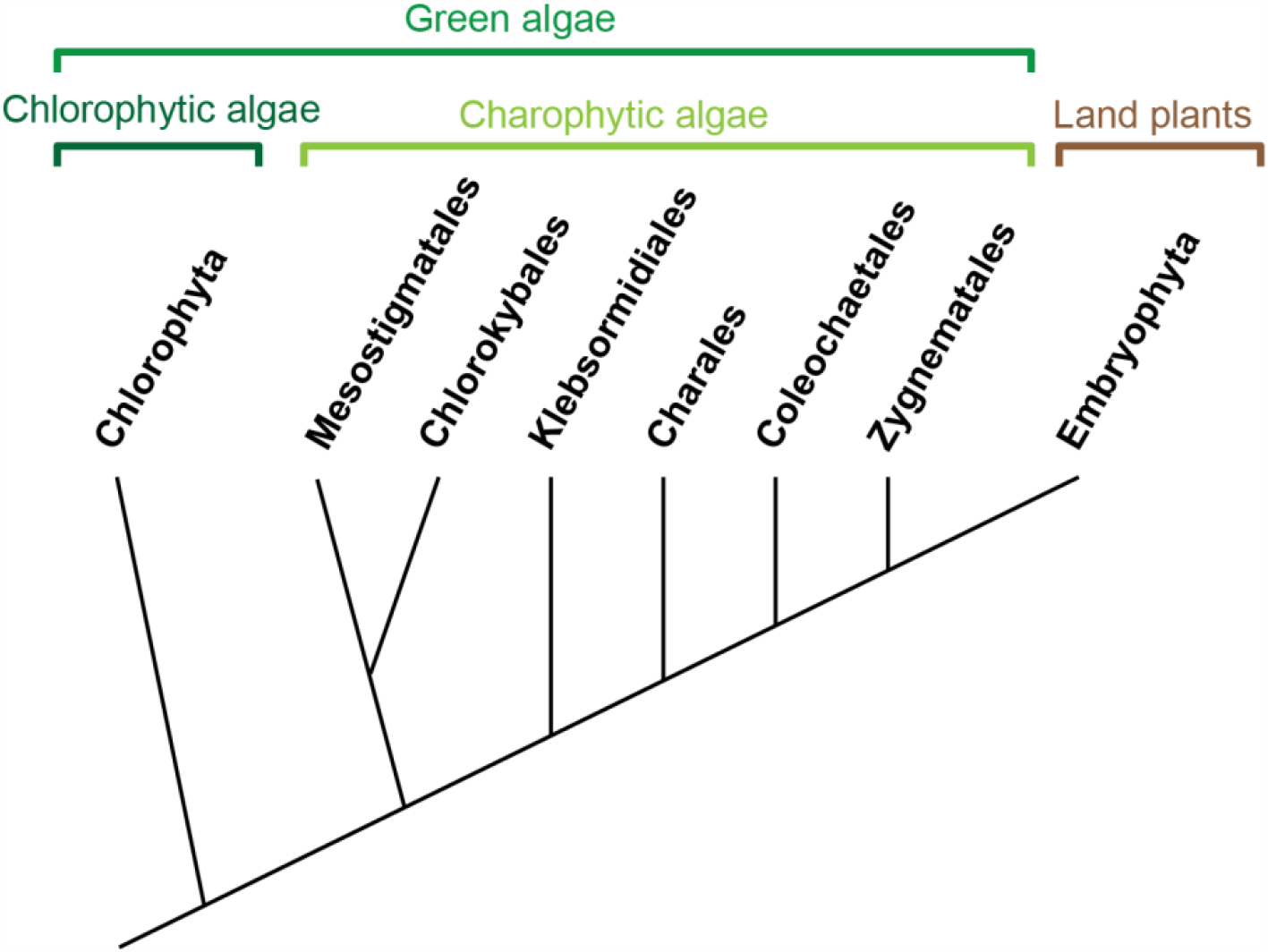
Evolutionary relationships of CGAs to other lineages in Viridiplantae. The monophyletic Chlorophytic green algae are sister to the polyphyletic Charophytic green algae. Species in the Zygnematales and the closest algal relatives to land plants (de Vries & Archibald, 2018).

**Supplemental Figure S2.**
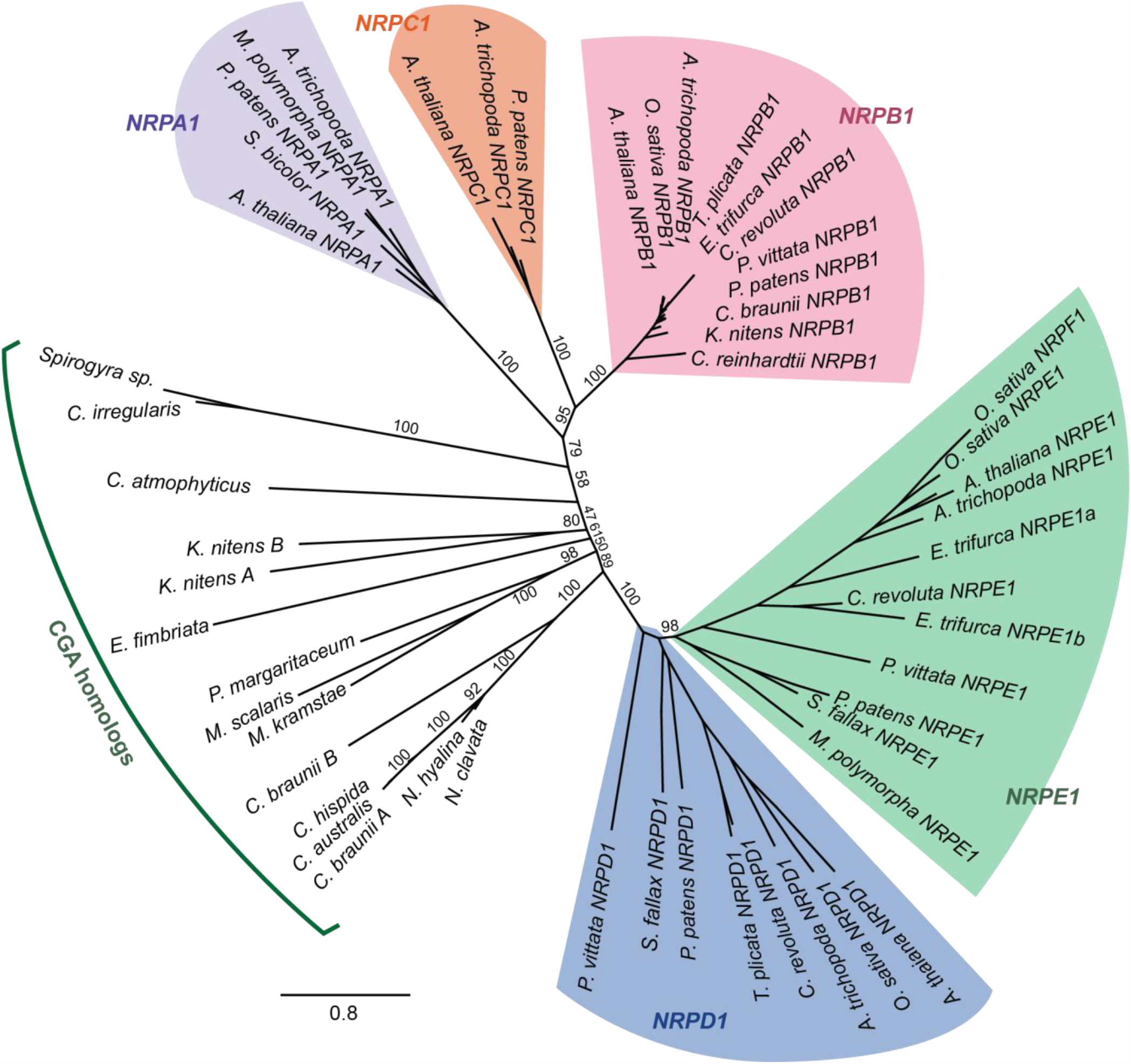
Phylogenetic relationship between the largest subunits of Pol I-V (NRPA1, NRPB1, NRPC1, NRPD1, and NRPE1) and CGA sequences shows that CGA NRPD/E1 homologs are a distinct group from other RNA polymerase largest subunits. Amino acid sequences were aligned with MAFFT v7.450 and stripped in positions where 50% of the taxa contained a gap. The tree was inferred by maximum likelihood.

**Supplemental Figure S3.**
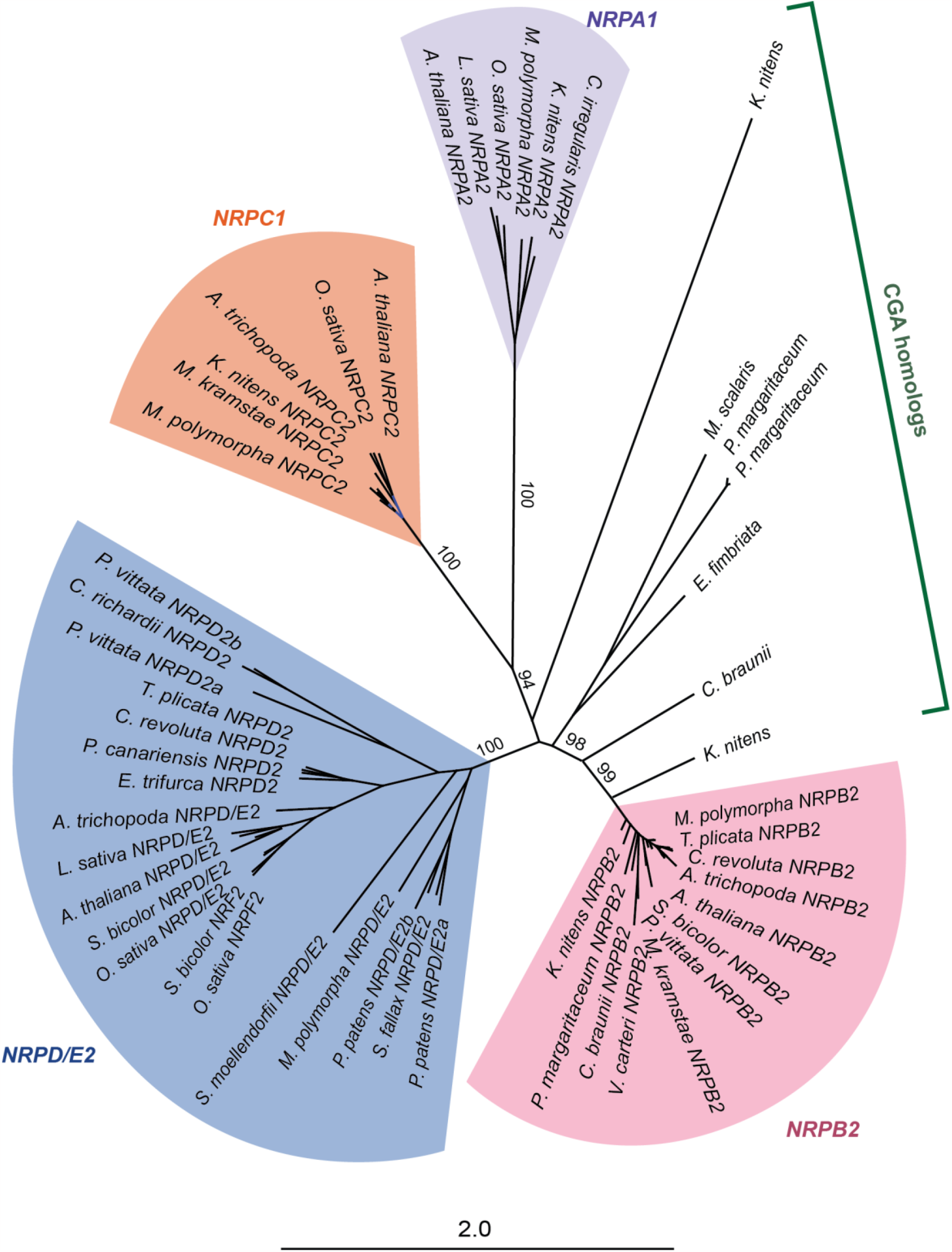
Phylogenetic relationship between the second subunits of Pol I-V (NRPA2, NRPB2, NRPC2, and NRPDE2) and CGA sequences shows that CGA NRPD/E2 homologs are a distinct group from other RNA polymerase largest subunits. Amino acid sequences were aligned with MAFFT v7.450 and stripped in positions where 50% of the taxa contained a gap. The tree was inferred by maximum likelihood.

**Supplemental Figure S4.**
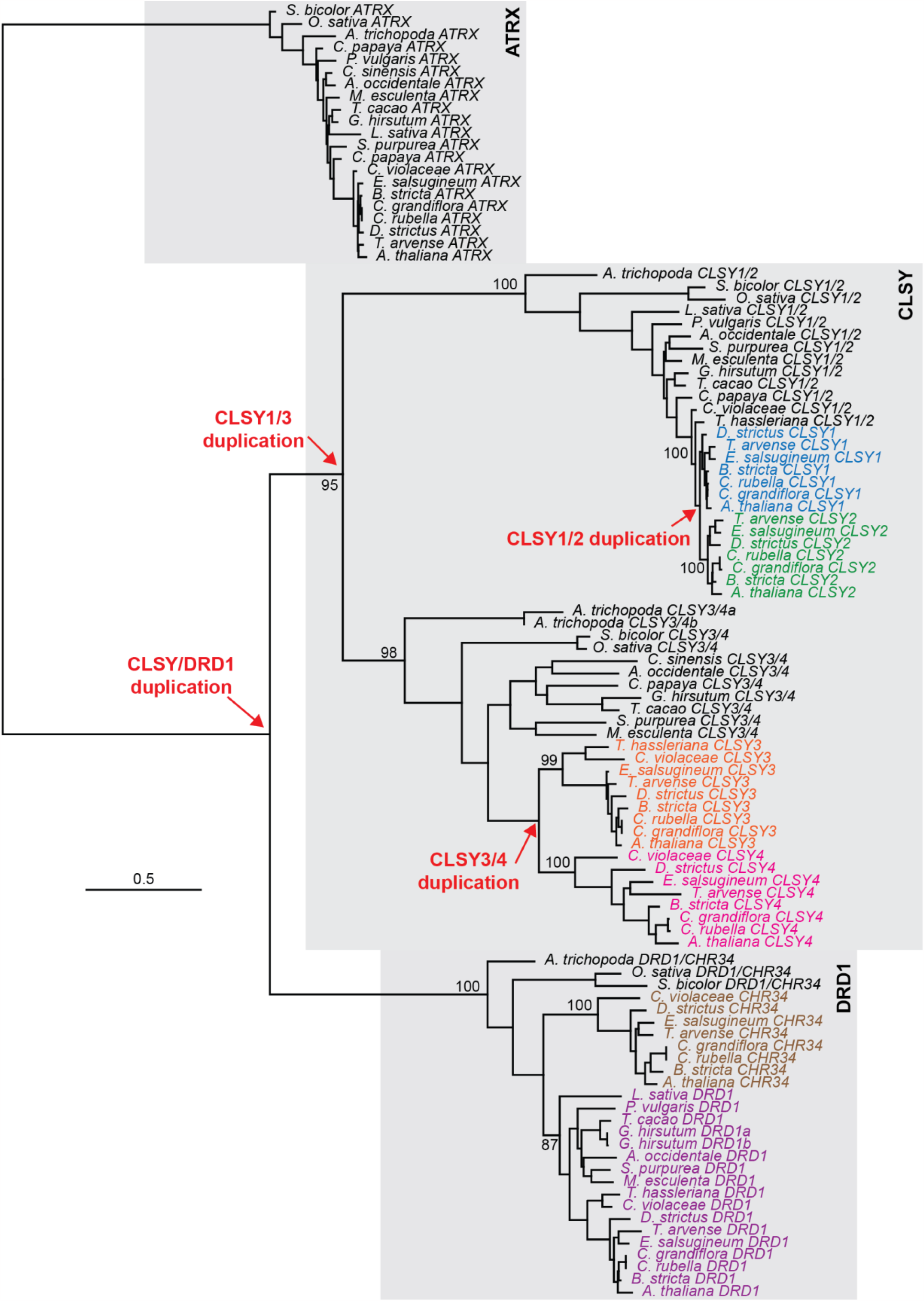
Duplication history of *CLSY* and *DRD1* clade. Amino acid sequences were aligned with MAFFT v7.450, the conserved SNF2 and helicase domains and intervening sequences were extracted, realigned, and stripped in positions where 50% of the taxa contained a gap. The tree was inferred by maximum likelihood and rooted on the *ATRX* outgroup. Deeper sampling of taxa in Brassicaceae and eudicots demonstrates that the duplication giving rise to *CLSY1* and *CLSY2* occurred at the emergence of Brassicaceae, as there are only single-copy *CLSY1/2* homologs in *T. hasseleriana* and *C. violaceae* (in the Cleomaceae family). In contrast, the duplication giving rise to *CLSY3* and *CLSY4* is slightly older, occurring sometime between the divergence of the Caricaceae (*C. papaya*) and Cleomaceae families within the Brassicales order.

## Notes

### Competing Interest Statement

The authors have declared no competing interest.

